# Features affecting Cas9-Induced Editing Efficiency and Patterns in Tomato: Evidence from a Large CRISPR Dataset

**DOI:** 10.64898/2026.01.06.696182

**Authors:** Amit Cucuy, Daniela Ben-Tov, Cathy Melamed-Bessudo, Arik Honig, Barry A. Cohen, Avraham A. Levy

## Abstract

CRISPR/Cas9 is a cornerstone of genome editing, yet the determinants of editing efficiency and DNA Double Strand Break (DSB) repair outcomes remain poorly understood, particularly in plants. To address this gap, we generated a dataset of 420 sgRNAs targeting promoters, exons, and introns of 137 genes in tomato protoplasts and quantified editing efficiencies together with ATAC-seq–derived chromatin accessibility and transcriptional states in the same cellular system. Editing efficiency was consistently higher at open chromatin sites and modestly elevated in promoters and introns relative to exons, whereas transcriptional activity did not measurably influence editing outcomes. Additionally, we identified a local genomic effect resulting in less variable editing between sgRNAs targeting the same compared to different genes. A distinct subset of sgRNAs achieved nearly complete editing, producing long deletions with extended microhomology tracts. These repair footprints closely parallel those observed for high-efficiency guides in human datasets, implicating conserved sequence-driven biases and a predominant role for microhomology-mediated end joining at these sites. Yet, widely used human-trained prediction models failed to rank sgRNA performance in plants, underscoring the limits of cross-species generalization.

This dataset defines how chromatin accessibility, genomic context, and intrinsic sequence characteristics shape Cas9 activity in plants, and provides a resource for improving guide design and advancing mechanistic understanding of plant DNA repair.

## Introduction

Over the last decade, a major effort has been made to collect data that would enable us to predict and understand what determines genome editing efficiency. This is a complex task due to the multiplicity of parameters that could affect the cleavage and the repair of CRISPR-Cas9-induced DNA double-strand breaks (DSBs). Large datasets have been generated particularly in mammalian cell culture systems, with studies examining up to 40,000 different single guide RNAs (sgRNAs) (Allen et al. 2019) and others analyzing several thousands (Leenay et al. 2019; Chakrabarti et al. 2019; Shen et al. 2018).

Such large-scale experiments are possible because individual cells can be independently transformed, allowing each cell to represent a separate editing event and enabling the capture of the full spectrum of repair outcomes. For example, Allen et al. (2019) recorded more than 10⁹ mutational outcomes across 40,000 guides. These and other studies have identified several factors influencing both repair outcomes and overall genome-editing efficiency. Early studies established that repair following Cas9-induced double-strand breaks is not random: consistent, sequence-dependent patterns of insertions and deletions emerge across experiments. Comparisons between different cell lines and delivery systems revealed that one of the most prominent determinants of repair-outcome distribution is the target-site sequence itself (van Overbeek et al. 2016; Chakrabarti et al. 2019).

On a broader scale, systematic differences have been observed between organisms. Although repair outcomes follow reproducible trends, they remain only partially predictable. Beyond sequence composition, multiple factors influence editing outcomes. For example, cross-species variation in DNA-repair machinery, as shown by Weiss et al. 2024), can alter the relative activity of repair pathways and thus the resulting mutation patterns. Epigenetic features also differ across cell types and tissues and have been shown to influence repair-pathway choice (Schep et al. 2021). In both plant and mammalian systems, regions of heterochromatin generally show lower mutagenesis than euchromatin (Weiss et al. 2024; Horlbeck et al. 2016; Schep et al. 2021), suggesting that chromatin context could be a key determinant of editing efficiency. Finally, transcriptional activity has been shown to affect both the likelihood and outcome of repair (Horlbeck et al. 2016).

Despite major progress, predictive models developed in mammalian systems still explain only part of the variability in editing outcomes. In fact, when applied to plants, their accuracy drops sharply, reflecting fundamental differences in repair determinants (Weiss et al. 2024; Slaman et al. 2023). This limited predictability underscores the need for plant-specific datasets that directly link sequence, chromatin, and transcriptional features to editing efficiency. Unlike mammalian systems, where diverse cell lines enable high-throughput analysis of editing outcomes, most plant genome editing studies rely on whole regenerated plants (Gao 2021). This vastly limits the quantity of editing outcomes that can be analyzed, as the regeneration step creates a bottleneck, with only a few early editing events propagated in the plantlet.

An additional experimental system for measuring genome editing efficiency in plants utilizes protoplasts, cell-wall-free cells that are released through enzymatic digestion. While protoplast-based systems have been used mainly as a proxy for evaluating sgRNA activity (Gao 2021), their unique properties also make them particularly well-suited for studying DNA repair dynamics in plants. In addition to their high transformation efficiency and suitability for large-scale experiments, protoplasts offer a key biological advantage: they do not divide in the absence of inducing factors and therefore do not undergo clonal propagation (Ben-Tov et al. 2024). As a result, there is no risk of clonal expansion driven by selection, and the ratio of different mutations remains stable, reflecting the true distribution of repair outcomes. This property allows editing efficiency to be measured directly, without biases introduced by growth or regeneration. Furthermore, the use of pre-assembled Cas9 ribonucleoprotein (RNP) complexes provides a transient and uniform delivery method that supports accurate assessment of both editing efficiency and repair diversity (Kim et al. 2014).

In this work, we used an experimental system that enables direct and comparable measurements of editing efficiency across hundreds of target sites within a single, uniform cellular environment, systematically measuring editing efficiency and repair outcomes for 420 CRISPR/Cas9 guides in tomato protoplasts. The dataset includes sgRNAs targeting promoter, exon, and introns of 137 genes, including major breeding and regulatory genes in edible tomato, *Solanum. Lycopersicum* cv. M82. We examine how gene features, genomic and epigenetic context and transcription influence editing. Using this data, we identify the effect of gene feature on editing efficiency, confirm the role of chromatin accessibility on editing efficiency and describe a unique group of high-editors, achieving nearly 100% editing within 48 hours, and significantly associated with microhomologies.

## Results

### Generating a Dataset of 420 CRISPR/Cas9 sgRNAs defined by Genomic Feature, Chromatin Accessibility, and Transcriptional State

To investigate the characteristics impacting genome editing efficiency and repair outcomes in tomato, we generated a dataset of CRISPR/Cas9 sgRNAs targeting promoter, first exon, and first intron of 137 genes along the tomato genome (Fig. 1A-B). The genes were selected from a list of important breeding genes published by Rothan et al. (Rothan et al. 2019), combined with chromatin modifiers, DNA repair, and housekeeping genes (Supplementary Data 1). Altogether, the dataset contains 420 unique sgRNAs (Fig. 1A-B, Supplementary Data1). The targets are evenly distributed amongst the 12 chromosomes, with an average of 34 targets per chromosome, primarily located in the subtelomeric regions (Fig. 1B-C). The majority of genes are targeted by three sgRNAs, with some as many as seven and as few as 1 (Fig. 1B, D).

**Figure 1.**
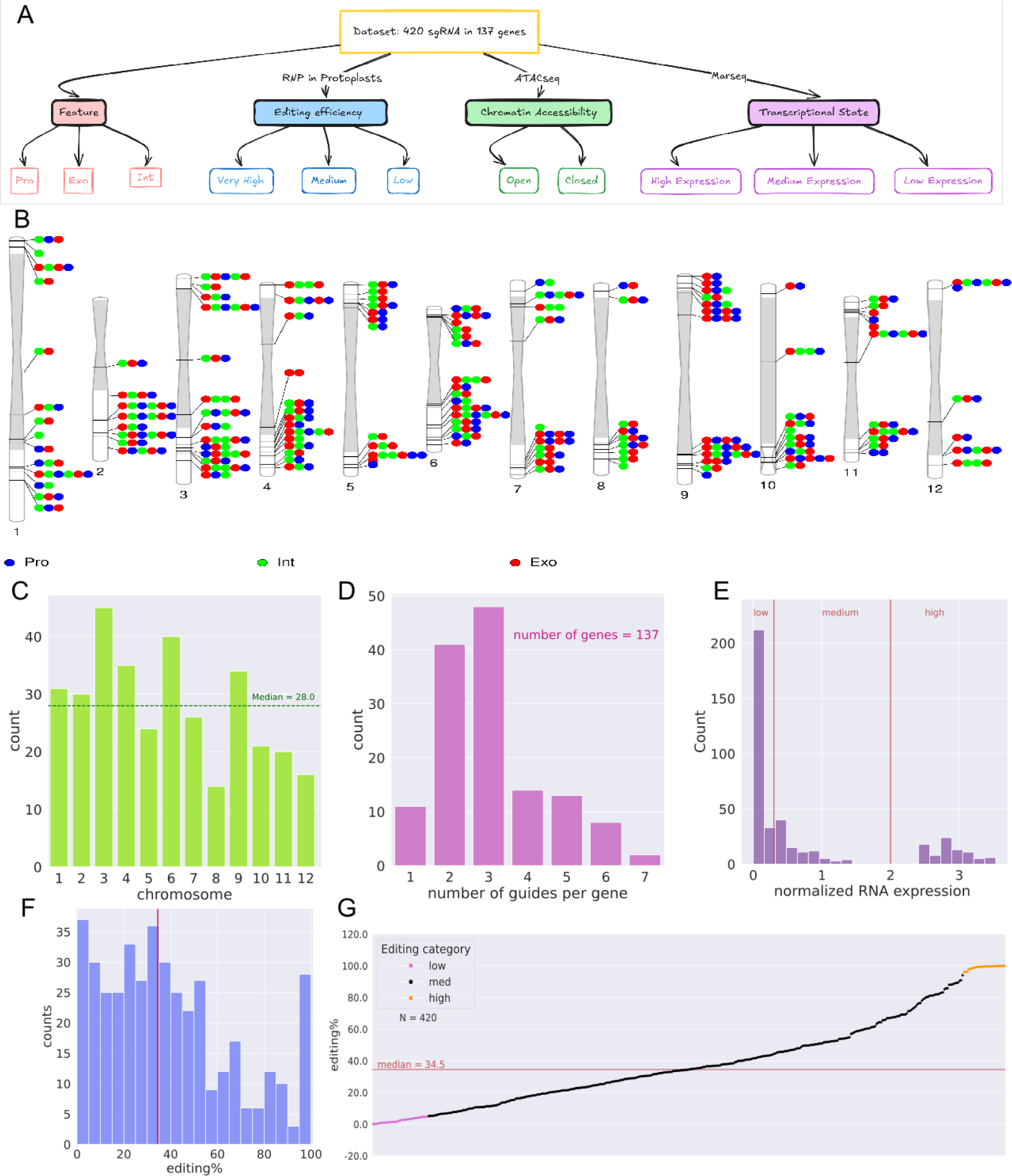
Experimental Design and Data Characteristics for CRISPR/Cas9 Guides and Editing Efficiency in Tomato Protoplasts. (A) Schematic of experimental design: 420 unique sgRNAs targeting 137 genes along the tomato genome. Each guide defined by feature (exon, intron, promoter), editing efficiency (low <5%, medium 5-95%, high >95%), chromatin accessibility (open vs. closed), and transcriptional state (low, medium, high). (B) Distribution of sgRNAs along the 12 tomato chromosomes. Each line represents a gene, and each dot corresponds to a unique guide, color-coded by genomic feature (Exon in red, intron in green, promotor in blue). The grey sections represent pericentromeric heterochromatin regions. (C) Number of guides per chromosome (the horizontal line representing the median). (D) Histogram of number of guides per gene (E) Histogram of normalized RNA-seq read count in log scale. Vertical red lines mark thresholds for categorical classification (low <2 reads, high >100 reads). (F) Histogram of editing efficiency (%) across 420 samples. The red vertical line indicates the median editing efficiency (34.5%). (G) Scatter plot of samples sorted by editing efficiency, with dots color-coded by editing categories (low <5%, high >95%). The red line represents the median

The transcriptional state of each gene was defined, through MARSeq (Keren-Shaul et al. 2019) conducted on the tomato protoplasts, as either low, medium or high expressing (Fig. 1A and E, Supplementary Data 1). In addition, ATAC-seq was conducted to determine the accessibility of the chromatin at each target site as either “open” or “closed” (Fig. 1A, Supplementary Data 1). Editing efficiency was determined 48 hours following PEG-mediated transformation of tomato protoplasts, through PCR Amplicon Sequencing (Fig. 1F-G). Since editing efficiency experiments were conducted in separate batches, 84 sgRNAs were selected and repeated in a normalization batch, used to account for batch effect (Fig. S1). From a total of 420, 94 sgRNA were transformed with preassembled CRISPR ribonucleoproteins more than once (between 2-5 biological replicates). Editing efficiencies and the mutational signature from these replicates were averaged to produce a single data point per guide, which was used in downstream analyses (see Supplementary Data 1).

The editing efficiency across the dataset shows a trimodal distribution, ranging from near 0% to nearly 100%, with a median of 34.5% (Fig. 1F–G). This resource provides a large, high-resolution dataset of sgRNAs with experimentally determined editing efficiency, categorically defined by genomic features, chromatin accessibility, and transcriptional state (Fig. 1A, Supplementary Data 1).

### Effect of Transcription, Chromatin Accessibility, and Genomic Feature on Editing Efficiency

Editing efficiency of CRISPR/Cas9 is affected by the context of the target sequence, its accessibility to the CRISPR RNP and to the repair machinery, and the processes working at the site (Schep et al. 2021; Verkuijl and Rots 2019; Přibylová et al. 2022; van Overbeek et al. 2016). Therefore, the genomic context defined by gene feature (exon, intron, promoter), chromatin accessibility, and transcriptional state can impact RNP cleavage and repair. To examine these factors, sgRNAs grouped by these characteristics were compared (Fig. 2). Chromatin accessibility significantly affects editing efficiency, with higher editing in open versus closed chromatin (Fig. 2A). There is a marginally significant effect of genomic feature (p=0.52 for the general feature effect), with higher editing in promoters compared to exons (Fig. 2B, p = 0.048). In contrast, transcriptional state had no detectable effect, with similar editing observed at targets in low-, medium-, and high-expressing genes (Fig. 2C).

**Figure 2.**
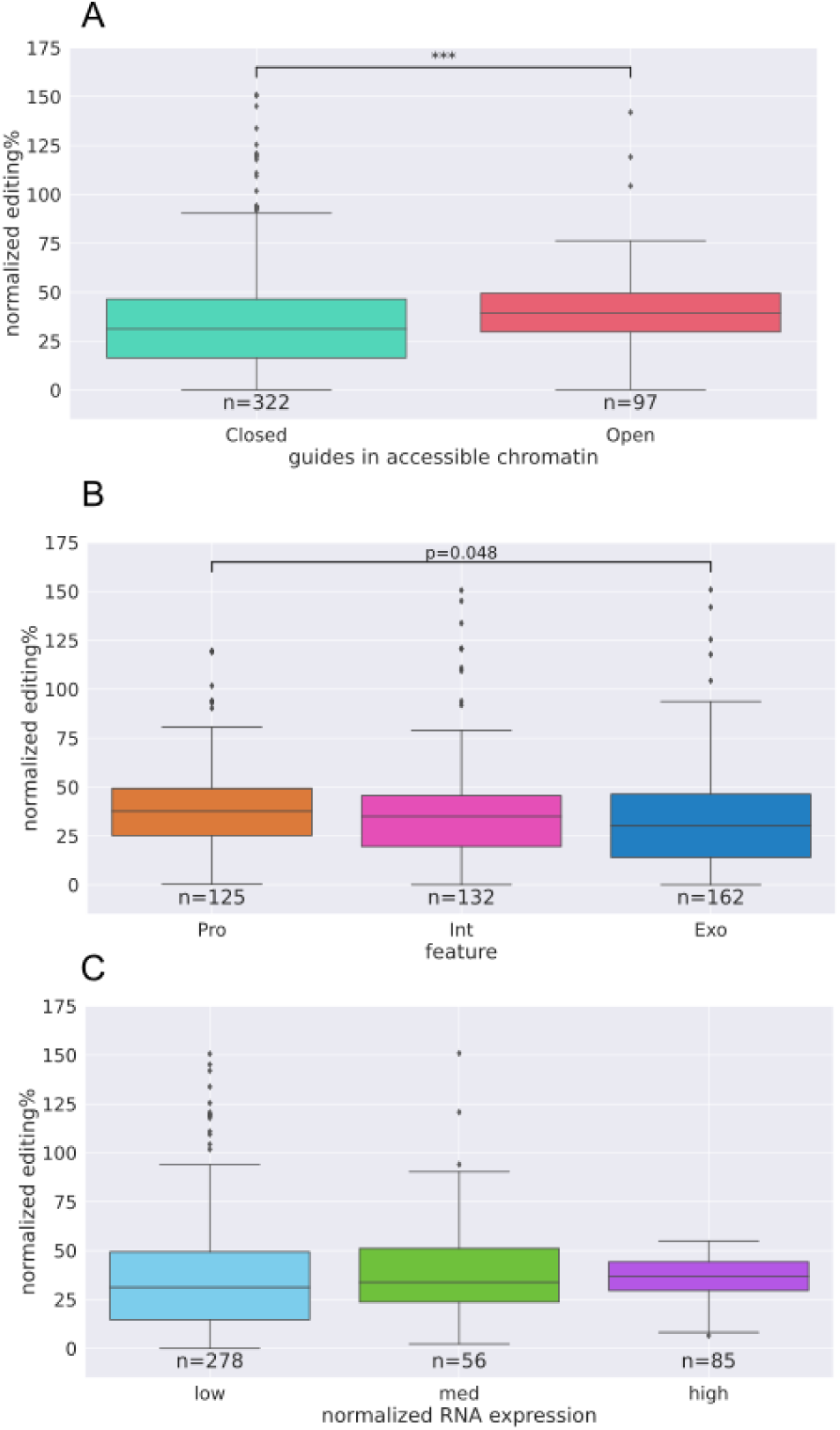
Effect of chromatin context, feature, and transcription on editing efficiency. A) Comparison of editing efficiency in open vs. closed chromatin (p < 0.005, Mann-Whitney test). B) Comparison of editing efficiency in different genomic features (promoter, intron, exon), the general effect of the feature was marginally significant (p=0.52, Kruskal-Wallis), the difference between Exon and promoterwas significant (p=0.048, Dunn’stest). (C) Comparison of editing efficiency in low, medium, and high expression targets.

### Higher variability between genes than within genes suggests local effects on editing efficiency

To further investigate the variability in CRISPR/Cas9 editing efficiency across our dataset, we tested whether guides targeting the same gene produce more consistent editing efficiencies than guides targeting different genes. We compared the standard deviation (SD) of editing efficiency within each gene to the global SD across all guides. This analysis reveals a lower within-gene SD (24.05) compared to the global SD (27.22; permutation test, p = 0.001; Fig. 3A and B), indicating that local genomic context influences editing outcomes. Nevertheless, significant differences may occur between nearby guides.

**Figure 3.**
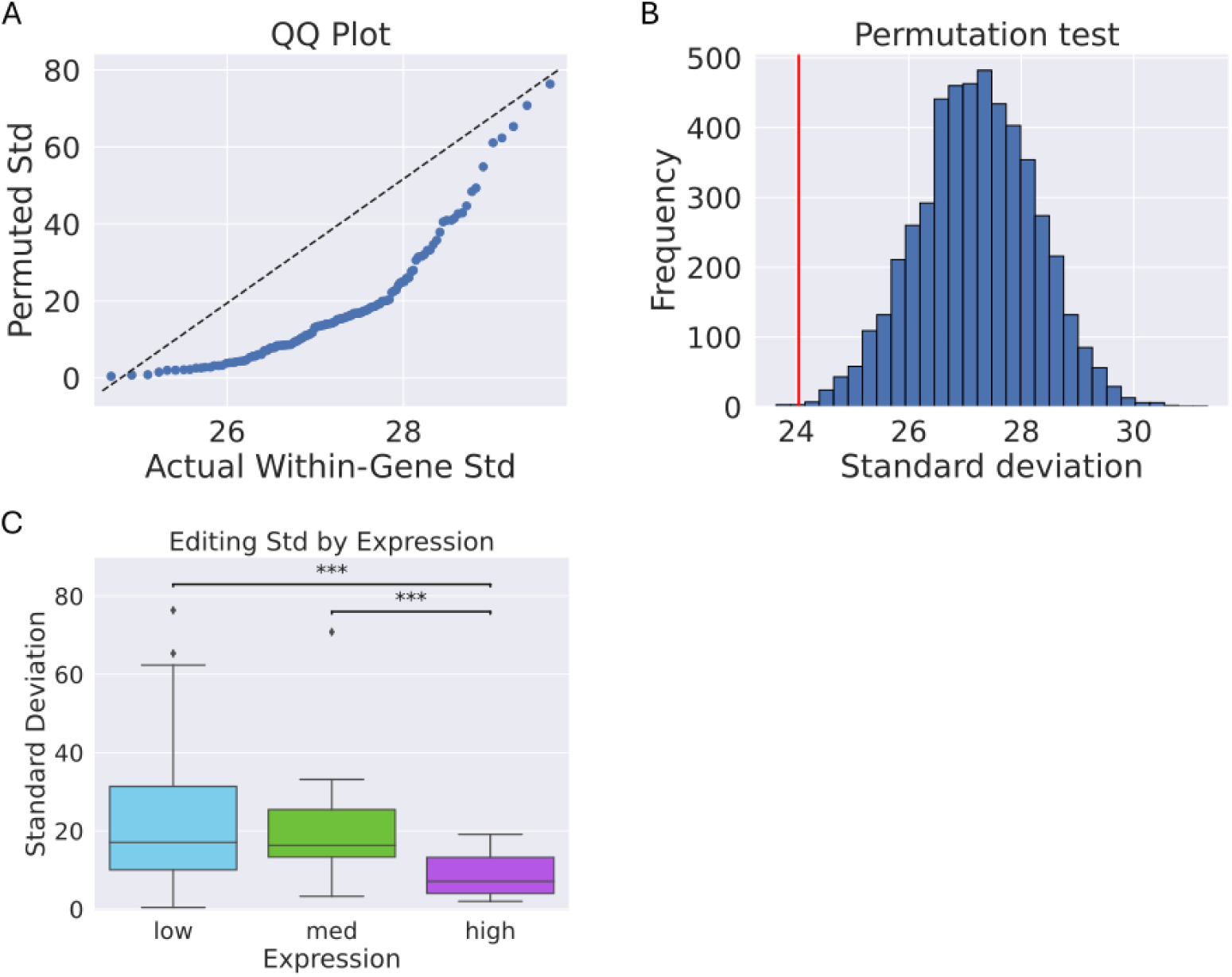
Local context influences editing efficiency within genes. **(A)** Quantile-quantile plot comparing the empirical distribution of observed within-gene standard deviations of editing efficiency to the empirical distribution obtained from permuted datasets; the dashed diagonal indicates equality between the two distributions. **(B)** Histogram of within-gene standard deviations from 10,000 permutations. The red line marks the observed value (24.05); the permuted mean was 27.22. The difference is statistically significant (p = 0.001). **(C)** Box plot showing within-gene variability in editing efficiency stratified by transcriptional activity. Highly expressed genes exhibit significantly lower within-gene variability than low-and medium-expression groups (p < 0.001).

We next asked whether transcriptional activity influences intra-gene variation. Highly expressed genes showed significantly lower within-gene variability compared to low- and medium-expression genes (Kruskal Wallis test, p < 0.001; Fig. 3C), suggesting that active transcription may stabilize editing outcomes across different target regions. Moreover, pairwise correlations of editing efficiencies between guides targeting distinct features of the same gene are modest but significant (r ≈ 0.25, p < 0.05), whereas shuffled controls show no correlation (r ≈ 0, p > 0.5; Fig. S2). Overall, although some genes exhibit substantial internal variation, guides targeting different positions within the same gene tend to show more similar editing efficiencies to each other than to guides targeting other genes, indicating that editing outcomes are partly constrained by broader local genomic context beyond the immediate target site.

### Repair footprint distributions differ by editor efficiency and chromatin context

To better understand the nature of CRISPR/Cas9-induced DNA repair, we analyzed the composition of insertion and deletion (indel) mutations, referred to as “repair footprints”. These footprints were normalized to each sample’s total editing efficiency to allow comparison across samples regardless of their absolute editing efficiency. Overall, deletions were the dominant mutation type, comprising 81.6% of all mutations. Insertions were relatively rare, except for +1 insertions, which accounted for 17.1% of all mutations and 94.0% of all insertions. Among all repair outcomes, the most common mutations were −1 and +1 indels, together accounting for 37.0% of total mutations. When combined with deletions up to −4 bp, this subset represents nearly half (47.9%) of all observed mutations.

When we compared repair footprints across editing efficiency groups, clear differences emerged. In low-efficiency guides, there was a strong overrepresentation of −1 deletions (Fig. 4A), while high-efficiency guides lacked both −1 and +1 peaks and instead showed a pronounced increase in longer deletions (Fig. 4C). The increase in average deletion length among high editors was statistically significant (p < 0.01; Fig. 4D), and was accompanied by a decrease in average insertion length. Insertions were almost completely absent in high-editing guides (Fig. 4E).

**Figure 4.**
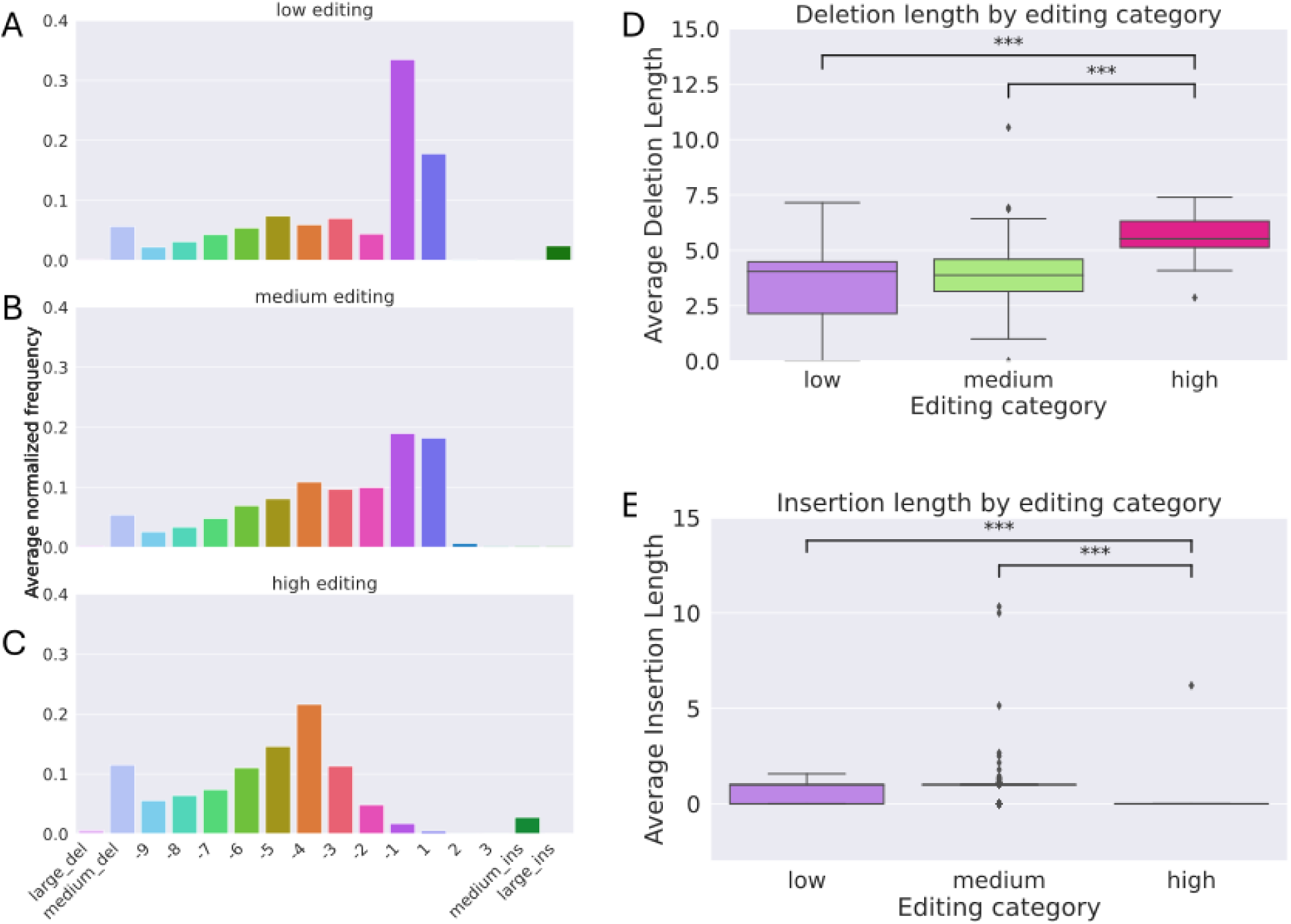
High-efficiency guides produce longer deletions and fewer insertions. **(A-C)** Average repair footprint composition across low-, medium-, and high-efficiency editing groups. The x-axis indicates individual indel sizes (e.g., -1, -2, +1, etc.), with additional bins for large deletions (>20 bp), medium deletions (10-20 bp), medium insertions (4-10 bp), and large insertions (> 10 bp). The y-axis shows the average normalized frequency of each outcome, calculated by dividing indel counts by the total editing efficiency for each guide. **(D)** Box plot of deletion lengths by editing efficiency group. High-efficiency guides exhibit significantly longer deletions than medium- and low-efficiency guides (p < 0.001, Kruskal-Wallis test; Dunn’s test for pairwise comparisons). **(E)** Box plot of insertion lengths by editing efficiency group. Insertions are significantly shorter in high-efficiency guides compared to the low and medium categories (p < 0.001, Kruskal-Wallis test; Dunn’s test for pairwise comparisons).

Chromatin context also influenced repair outcomes: deletions in closed chromatin regions are longer on average than those in open regions (Fig. 5C, p < 0.05). However, when distributions are stratified by both editing efficiency and chromatin accessibility, editing category (low, medium, high) remains the stronger predictor of footprint composition (Supplementary Fig. 3). In contrast, transcriptional state and genomic feature have no observable effect on repair footprint distribution (Supplementary Fig. 4). These findings suggest that highly efficient editing may engage different repair mechanisms or reflect biases toward more extensive end resection.

**Figure 5.**
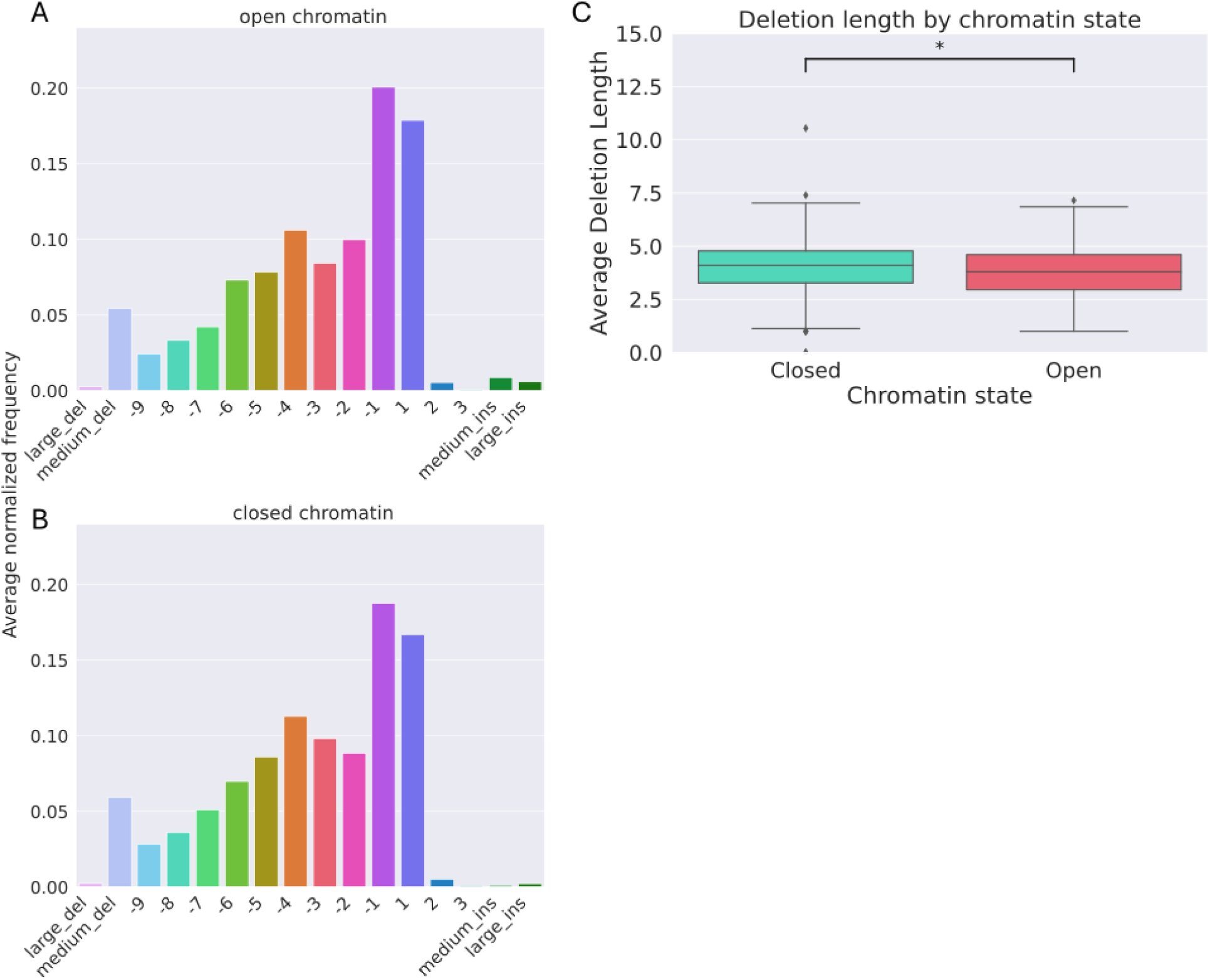
Deletion length is influenced by chromatin context. **(A-B)** Histograms showing the average repair footprint composition in open (A) and closed (B) chromatin regions. The x-axis represents InDel length categories, and the y-axis shows the average normalized frequency. **(C)** Box plot comparing average deletion lengths between open and closed chromatin. Deletions are significantly longer in closed chromatin (p < 0.001, Mann-Whitney U test).

### Microhomology-associated deletions are enriched in high-efficiency guides

Microhomology-associated deletions were defined as those where one breakpoint shares at least two consecutive bases with the sequence flanking the opposite end of the deletion. Analysis of deletion frequencies revealed no significant differences in the prevalence of these events across chromatin states (p = 0.125), transcriptional activity (p = 0.54), or genomic features (p = 0.99). However, high-efficiency guides exhibited a significant enrichment of microhomology-associated deletions compared to medium- and low-efficiency guides (p = 0.015; Fig. 6A–D). In addition, the average length of microhomology tracts was significantly longer in high-efficiency guides than in other groups (p = 4.69 × 10⁻⁵; Fig. 6H). No significant differences were observed when comparing chromatin state (Fig. 6E), transcriptional level (Fig. 6F) and genome features (Fig. 6G).

**Figure 6.**
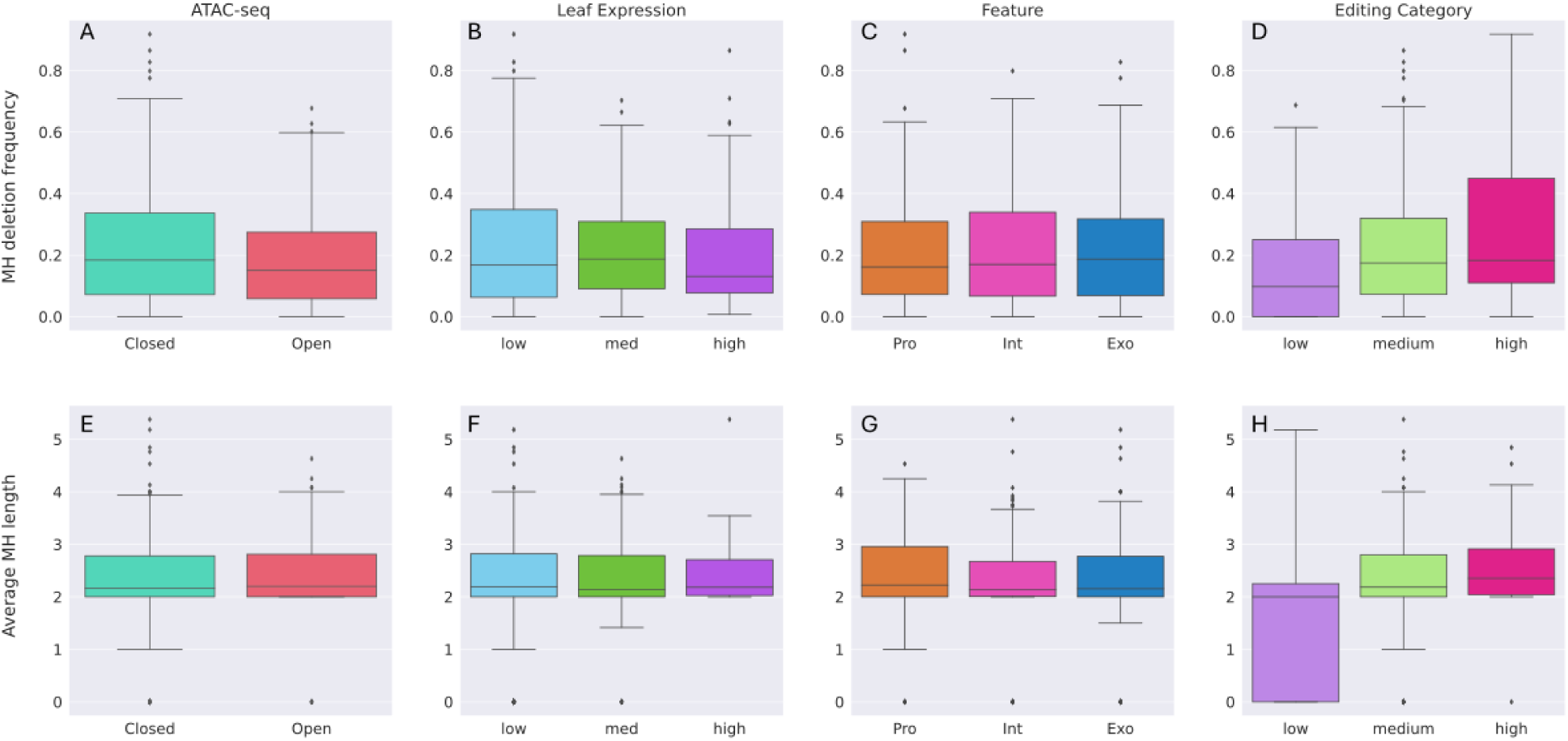
Microhomology-associated deletions are enriched in high-efficiency guides. **(A-D)** Box plots showing the frequency of microhomology (MH)-associated deletions across genomic and epigenetic categories. **(A)** MH deletion frequency in open vs. closed chromatin. Closed chromatin shows a non-significant increase in MH deletions (p = 0.125, Mann-Whitney U test). **(B)** MH deletion frequency across transcriptional states (low, medium, high); no significant differences observed (p = 0.54, Kruskal-Wallis test). **(C)** MH deletion frequency by genomic feature (exon, intron, promoter); no significant differences observed (p = 0.99, Kruskal-Wallis test). (D) MH deletion frequency by editing efficiency group (low, medium, high); high-efficiency guides show a significant enrichment compared to other groups (p = 0.015, Kruskal-Wallis test). **(E-H)** Box plots showing the length of microhomology tracts associated with deletions. **(E)** MH tract length in open vs. closed chromatin; no significant difference (p = 0.57, Mann-Whitney U test). **(F) MH** tract length across transcriptional states; no significant difference (p = 0.66, Kruskal-Wallis test). (G) MH tract length by genomic feature; no significant difference (p = 0.37, Kruskal-Wallis test). **(H)** MH tract length by editing efficiency group; significantly longer tracts observed in high-efficiency guides (p =4.69 × 10^-5^, Kruskal-Wallis test).

To explore possible sequence biases that may underlie this pattern, guides with >30% microhomology-associated deletions were compared to those below this threshold. Sequence-logo analysis revealed that high-microhomology guides were enriched for A/T bases adjacent to the break site, a bias absent in low-microhomology guides (Fig. 7). These findings suggest that microhomology-mediated end joining (MMEJ) contributes disproportionately to repair at high-efficiency sites, potentially facilitated by local A/T-rich sequence contexts.

**Figure 7.**
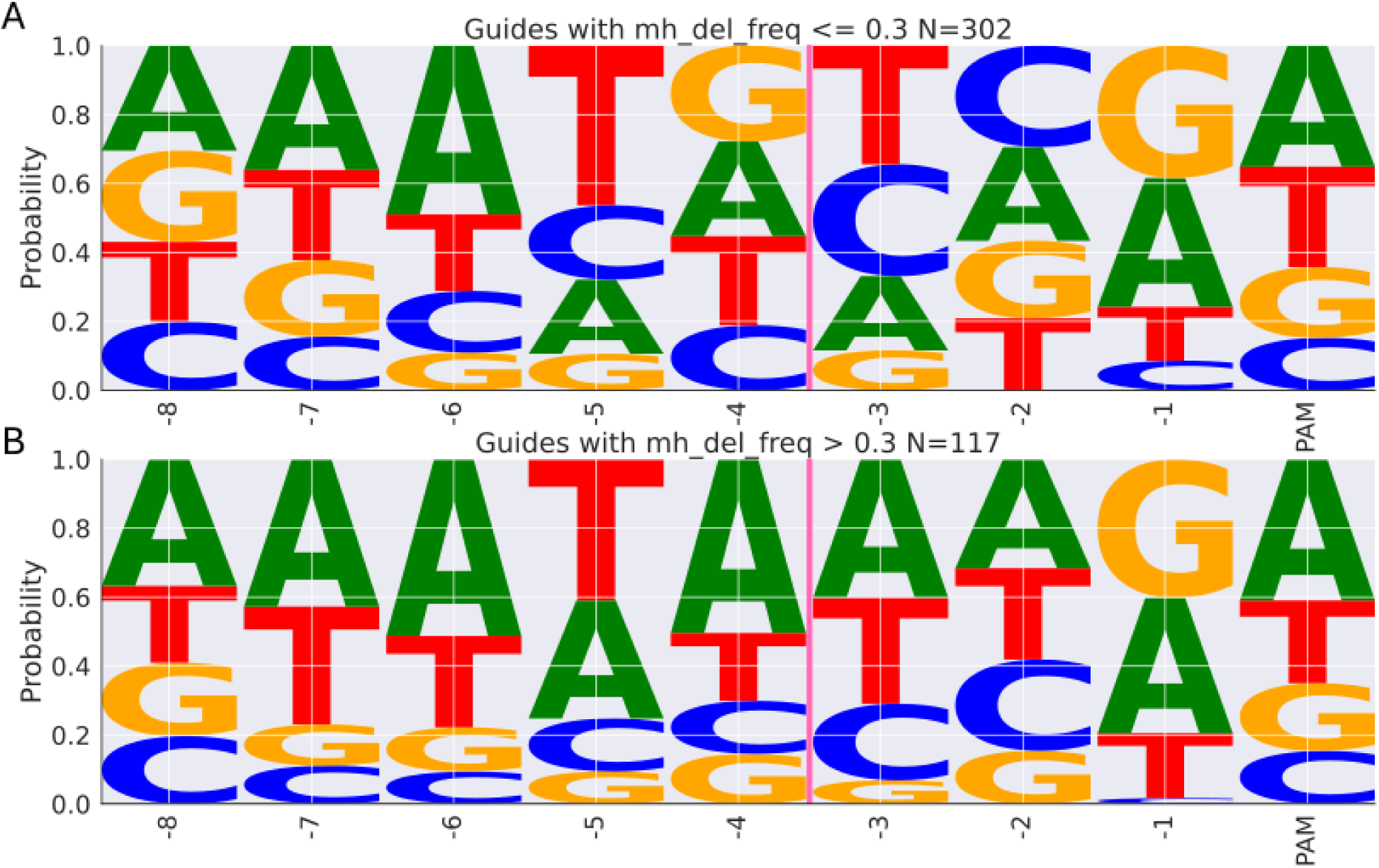
A/T enrichment in sgRNAs with high microhomology-associated deletion frequency. Sequence logos showing nucleotide composition of sgRNA target sites stratified by the frequency of microhomology-associated deletions. The x-axis represents position relative to the PAM sequence, with the first base of the PAM included. The pink vertical line marks the predicted Cas9 cleavage site. The y-axis shows the normalized frequency of each nucleotide at each position. **(A)** Sequence logo for sgRNAs with <30% microhomology-associated deletions (N = 394), showing relatively balanced base composition. **(B)** Sequence logo for sgRNAs with >30% microhomology-associated deletions (N = 147), showing strong enrichment of A/T bases near the PAM-proximal region.

### Sequence characteristics of high-efficiency guides are conserved between humans and tomatoes

To test whether sequence-driven repair biases are conserved across species, we compared our tomato protoplast repair footprints to a large-scale human dataset from primary T cells targeting 1,656 genomic sites (Leenay dataset;(Leenay et al. 2019)). In this human dataset, low-efficiency guides predominantly generate −1 deletions, while high-efficiency guides produce longer deletions with fewer insertions, mirroring the patterns observed in tomato (Fig. 8 A-E). These parallels suggest that intrinsic DNA sequence features similarly shape repair outcomes across species.

**Figure 8.**
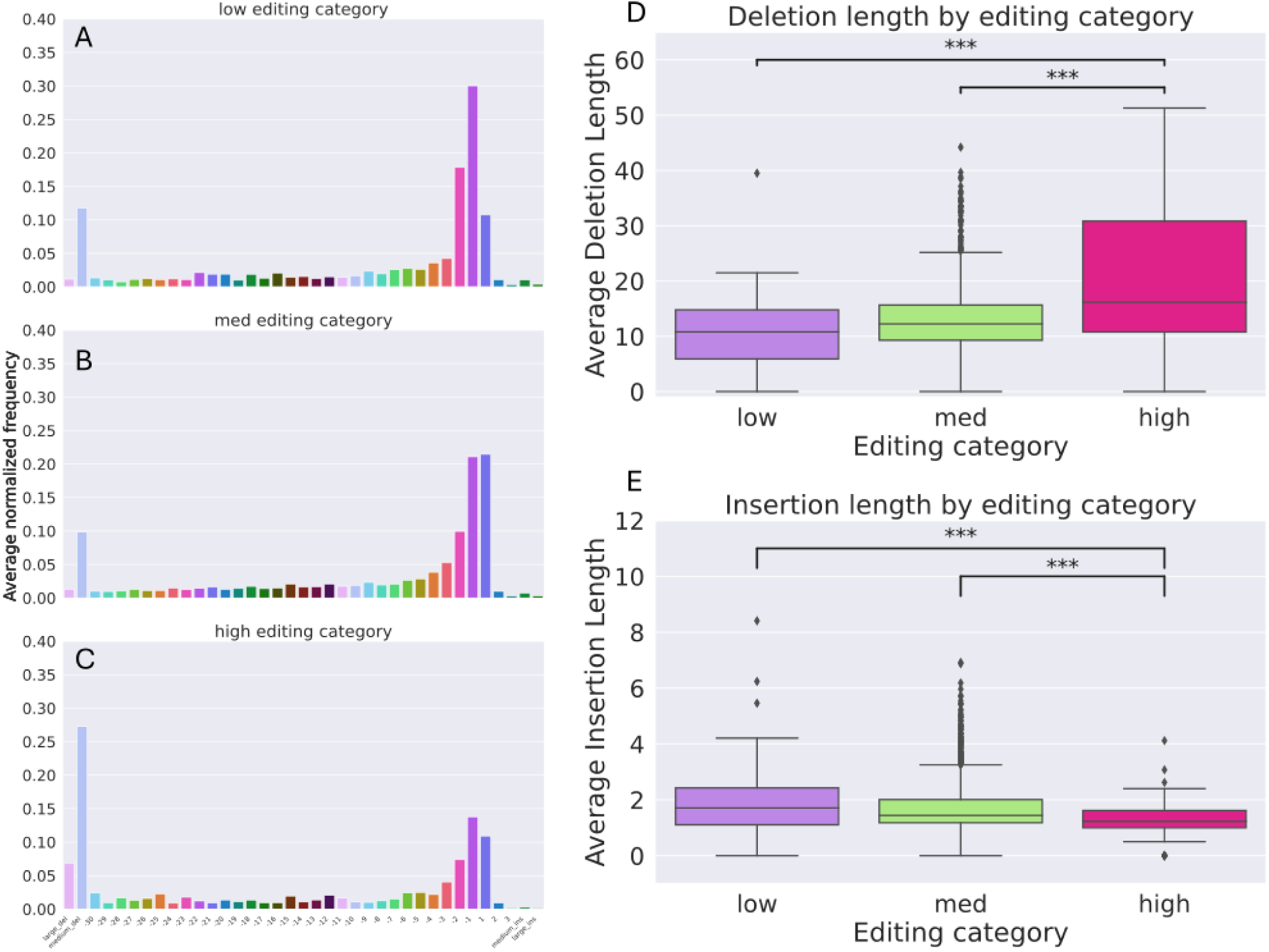
High-efficiency guides in the Leenay human dataset produce longer deletions and fewer insertions. **(A-C)** Average repair footprint composition for low-, medium-, and high-efficiency editing categories in the Leenay dataset (Leenay et al. 2019). The x-axis indicates indel length, and the y-axis shows average normalized frequency per guide. **(D)** Box plot of deletion lengths across editing efficiency groups. High-efficiency guides exhibit significantly longer deletions than medium- and low-efficiency guides (p < 0.001, Kruskal-Wallis test; Dunn’s test for pairwise comparisons).**(E)** Box plot of insertion lengths across editing efficiency groups. Insertions are significantly shorter in high-efficiency guides compared to low and medium categories (p < 0.001, Kruskal-Wallis test; Dunn’s test for pairwise comparisons).

We also analyzed the CRISPRon dataset, which provides on-target activity measurements for 10,592 SpCas9 sgRNAs in human cells (Xiang et al. 2021), to evaluate the cross-species applicability of predictive scoring models. We applied the Azimuth algorithm (Doench et al. 2016), a widely used machine learning model trained on human data, to both our tomato guides and the two human datasets. Azimuth scores, ranging from 0 to 100, showed a similar distribution across species (Fig. 9A). However, there was no significant correlation between Azimuth-predicted activity scores and measured editing efficiency in either dataset. When guides were stratified into low, medium, high, and very-high Azimuth bins, the human data showed significant differences in editing efficiency across bins, whereas tomato guides followed the same directional trend, but the difference was not statistically significant (Fig. 9B). These findings indicate that, while sequence context exerts a conserved influence on CRISPR repair footprints, prediction algorithms may require species-specific calibration to accurately rank sgRNA performance.

**Figure 9.**
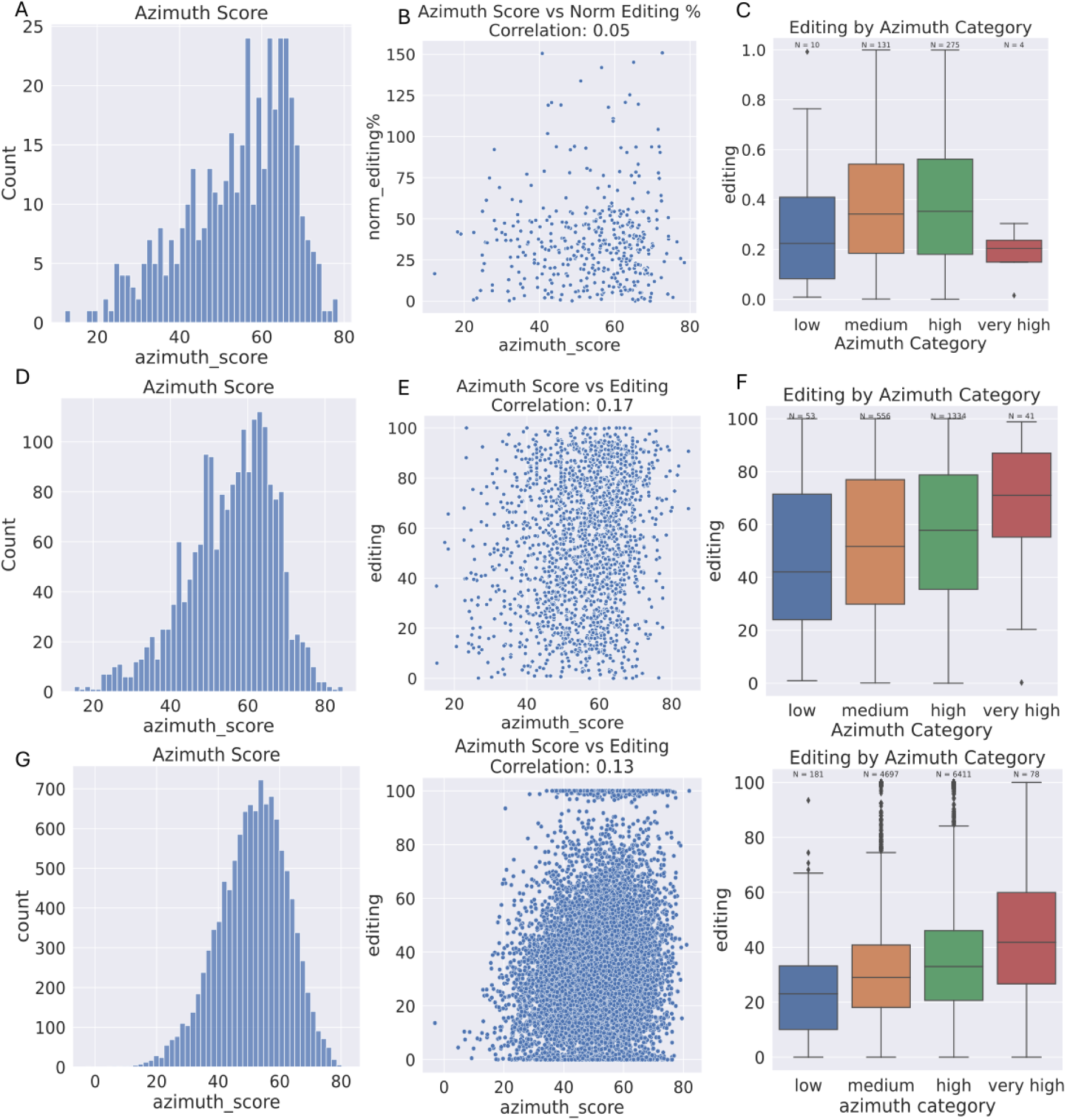
Evaluation of Azimuth score predictions versus measured editing efficiencies across tomato and human datasets. **(A-C)** Tomato protoplast dataset: **(A)** Histogram of Azimuth scores for tomato sgRNAs. **(B)** Scatter plot of Azimuth score versus normalized editing efficiency; no significant correlation observed (r = 0.05, p > 0.05). **(C)** Box plot of editing efficiencies stratified by Azimuth score category(low: N = 10; medium: N = 131; high: N = 275; very high: N = 4); no significant differences between groups (Kruskal-Wallis test, p > 0.05). (D-F) Leenay dataset (Human): **(D)** Histogram of Azimuth scores for human sgRNAs. **(E)** Scatter plot of Azimuth score versus editing efficiency; weak correlation (r = 0.17, p > 0.05). **(F)** Box plot of editing efficiencies by Azimuth score category; significant differences across groups (Kruskal-Wallistest, p < 0.001). (G-1) CRISPRon dataset (Human): **(G)** Histogram of Azimuth scores for sgRNAs. **(H)** Scatter plot of Azimuth score versus editing efficiency; weak correlation (r = 0.13, p > 0.05). **(I)** Box plot of editing efficiencies by Azimuth score category; significant differences across groups (Kruskal-Wallis test, p < 0.001).

## Discussion

### Chromatin and Genomic Features Shape Editing Outcomes

In this work, we generated several novel datasets for tomato, providing a new public resources for the amelioration of this important crop through genome editing: 1- A comprehensive list of CRISPR/Cas9 sgRNA sequences promoters, introns and exons of essential breeding genes expanded from Rothan et al. 2019; 2- a large dataset of editing efficiency measurements and repair footprints obtained for distinct sgRNAs at targeted loci, a rarity in plants; and 3- new *Solanum Lycopersicum* cv. M82 protoplast ATAC-seq and MARS-seq data. Taken together, these datasets enabled us to characterize the endogenous factors constraining genome editing efficiency and outcome as discussed below.

Our results showed that editing efficiency was higher in open chromatin, showed a marginally significant increase in promoters compared with exons, and was unaffected by transcriptional state, adding support to previous reports that chromatin accessibility influences CRISPR/Cas9 double-strand break induction and repair outcomes (Schep et al. 2021; Weiss et al. 2022; Liu et al. 2019), and that incorporating chromatin accessibility and other epigenomic layers improves predictive modeling of editing efficiency (Ito et al. 2024). Although previous work in mammalian systems linked transcriptional activity to increased editing efficiency and altered repair patterns (Aymard et al. 2014; Horlbeck et al. 2016), our results show no detectible effect in plant protoplasts, suggesting, potentially, that chromatin accessibility rather than transcriptional activity is the dominant determinant of Cas9-induced mutagenesis in this system. (Fig. 2). Despite this, on a finer scale, guides targeting promoters and introns lead to a higher percentage of indel accumulation than exons guides (Fig. 2B). This might be due to a higher repair fidelity in exons than in non-coding regions. Alternatively, this might result from a more efficient DSB induction in non-coding than in coding region. The recent findings of Ben-Tov et al. (2024), on a small set of genes, suggest that repair efficiency rather than DSB induction is more likely to explain editing efficiency. Similarly, Monroe et al. 2022) found a reduction in mutation accumulation in genes compared to intragenic regions and suggested that this is due to increased repair fidelity (Quiroz et al. 2024). These results suggest a variable repair fidelity across the genome, leading to a more constrained occurrence of mutations in evolutionarily important regions. How the repair machinery distinguishes between coding and non-coding DNA is a fascinating question for future research.

We also observed that local genomic context plays a role in determining editing efficiency. Guides targeting different sites within the same gene produced different levels of editing, yet this variability was smaller than that observed across different genes (Fig. 3). This indicates that constraints imposed by the local context define editing outcomes at a finer scale. Notably, although transcriptional state did not have a strong global effect, highly expressed genes tended to show reduced variability between guides, pointing to a stabilizing influence within genes (Fig. 3). Potentially, factors such as position within the nucleus, local chromatin architecture, and others limit the accessibility to repair machinery or DSB induction. Further studies can help define these constraints. In addition, we consistently observed enrichment of A/T bases in deleted sequences, including those not associated with microhomology, suggesting that intrinsic sequence biases contribute to repair outcomes (Supplementary Fig. 5), similar to what was previously found in human cell lines (Chakrabarti et al. 2019; van Overbeek et al. 2016).

### A Distinct Subset of High-Efficiency Editors

Within our dataset, we identified a unique subset of CRISPR guides that achieved near-complete editing efficiency (∼100% indels) within 48 hours (Fig. 1 G and F). These guides included those targeting all three feature types and chromatin contexts. However, they share some intrinsic properties consistent with repair via MMEJ, characterized by larger deletions, a near absence of insertions, and longer stretches of microhomology at deletion junctions (Fig. 4, 6D and H). This repair pathway, which was found to produce extended deletions in comparison to NHEJ after Cas9 DSB induction (van Overbeek et al. 2016), is considered highly error-prone, potentially providing a plausible explanation for the unusually high efficiency observed (Sfeir et al. 2024). Supporting this interpretation, sequence-logo analysis revealed that high-microhomology guides were enriched for A/T bases adjacent to the break site (Fig. 7). A similar connection between sequence composition and the likelihood of MMEJ has been reported previously: flexible A/T-rich loops promote synthesis-dependent MMEJ, whereas rigid poly(A) loops inhibit it (Hanscom et al. 2022).

While guides of such high efficiency are not well-characterized in plants, we identified a comparable class of high-efficiency guides in human CRISPR datasets (Fig. 8), indicating that the sequence-driven determinants of repair pathway choice may be conserved across species. Together, these findings suggest that A/T-rich sequence contexts may facilitate microhomology annealing, reinforcing MMEJ as the predominant pathway at high-efficiency sites across eukaryotes. Likely, these targets are both characterized by highly efficient DSB induction coupled with primarily error-prone repair, presumably through MMEJ. Further studies characterizing targets such as these, including in mutant backgrounds, can help further elucidate the mechanisms involved. This conservation suggests that MMEJ, an intrinsically error-prone repair pathway mediated by Pol θ and conserved across eukaryotes (Sfeir et al. 2024), produces characteristic mutational signatures, deletions joined through short microhomologies and sequence-dependent biases, that are similar in plants and animals. A deeper understanding of the factors governing pathway choice between MMEJ and NHEJ could help improve the efficiency and predictability of genome-editing outcomes.

### Conservation and Limits of Predictability

While our results reveal conserved sequence-driven biases in repair outcomes, they also highlight the limited predictability of editing efficiency across species. On one hand, both our plant dataset and human datasets showed parallel repair footprints: low-efficiency guides predominantly produced −1 deletions, whereas high-efficiency guides yielded longer deletions with minimal insertions (Fig. 5 and 8). This similarity highlights both the consistent impact of intrinsic DNA sequence characteristics on the repair outcome and the similarity and conservation of repair pathways across higher eukaryotes. On the other hand, Consistent with previous reports, we confirm that cross-species predictability remains limited, as models like Azimuth fail to apply to plant data (Fig. 9). (Weiss et al. 2024; Slaman et al. 2023). This gap emphasizes that predictive accuracy is constrained by additional species-specific factors.

Altogether, our findings contribute to an emerging model in which editing outcomes are likely shaped by multiple layers of influence, each operating at a different scale. We report on the sequence level, namely the A/T sequences and repeats; the coding versus-non coding regions effect; the relative similarity of guides proximity (hundreds or thousands of bp apart) and the chromatin accessibility. Additional factors such as tissue type and cell cycle stage were shown to act across the genome or specific chromosomes in mammalian cells (Schep et al. 2021; Lin et al. 2014). Our system, is derived from mesophyll cells and remains entirely in G1 (Ben-Tov et al. 2024), limiting the scope of the results. Further study of protoplasts derived from different tissues and from cycling cells, together with the knockdown of specific DNA repair genes may expand our understanding of the effect of these factors in plants. These layers collectively define and constrain the probabilistic landscape of repair outcomes.

## Methods

1. Plant material- *M82 tomato* (*Solanum lycopersicum* cv. M82) seeds were surface sterilized using a wet sterilization protocol. Seeds were first washed in 70% ethanol for 30 seconds, followed by a rinse with sterile water. They were then incubated in a sterilization solution containing 2% sodium hypochlorite (NaOCl) and 0.1% Tween 20 for 15 minutes with gentle shaking. Following sterilization, seeds were rinsed three times with sterile water and sown on Nitsch medium in Magenta boxes. Plants were grown under long-day conditions (16 h light/8 h dark) at 23 °C.
2. Protoplasts isolation Protoplasts were isolated as previously described (Ben-Tov et al. 2024). Briefly, the first true leaves from 16–21-day-old *M82* seedlings (approximately 24 seedlings) were excised and cut into thin strips. The tissue was incubated in a Petri dish together with 15 mL of enzyme solution (Cellulase and Macerozyme), in the dark with gentle shaking (25 RPM) for 14–16 hours. Following digestion, the released protoplasts were filtered through a 100 µm nylon mesh and washed with W5 solution. Intact protoplasts were separated using a 23% sucrose gradient and further washed with W5. The final protoplast suspension was adjusted to a working concentration of 1 × 10⁶ cells/mL in MMG solution.
3. Cas9 ribonucleoprotein (RNP) complexes were prepared and transformed into protoplasts as previously described (Ben-Tov et al. 2024) with minor modifications. Briefly, 10 µg purified SpCas9 protein and 20 µg duplexed sgRNA (crRNA + tracrRNA; IDT) were mixed in NEB Buffer 3.1 (20 µL total) and incubated for 15 min at room temperature to assemble the RNP. crRNAs were designed to target the first exon and intron (when possible), and the promoter region (1 kb upstream of the TSS). The RNP mixture (20 µL) was added to 200,000 protoplasts in 200 µL MMG solution, followed by gentle mixing with 200 µL 40% PEG 4000 solution. After 20 min incubation in the dark, samples were diluted with W5 solution, incubated for 15 min, pelleted (450 × g), resuspended in 1 mL WI buffer, and incubated at 22 °C in the dark. After 48 h, protoplasts were pelleted, flash-frozen in liquid nitrogen, and genomic DNA extracted using the NucleoSpin Plant II kit (Macherey-Nagel) with minor modifications: 600 µL PL1 buffer + 10 µL RNase A, 1 h incubation at 65 °C, followed by column purification and elution in 50 µL buffer.
4. Library preparation and sequencing Libraries were prepared using a two-step PCR amplification protocol. In the first PCR step, genomic DNA was amplified using KAPA HiFi HotStart ReadyMix (Roche) and target-specific primers (supplementary table 1). These primers contained the target-specific sequence and a tail with part of the P7 or P5 Illumina sequence. The first PCR products were purified using SPRI beads then 2 µL of the purified first-round product, KAPA HiFi, and Illumina enrichment primers were used for the second PCR step Final libraries were purified using a two-step SPRI bead cleanup: first with 0.7× bead volume to remove large fragments, followed by 1.2×–1.5× to select the desired amplicon size. libraries were sequenced on an Illumina Novaseq platform using 2 × 150 bp paired end reads.
5. MARS-seq MARS-seq libraries were prepared as described in Keren-Shaul et al. (Keren-Shaul et al. 2019). The RNA was extracted from 200,000 cells of freshly isolated protoplasts and from a single leaf from the same plants used for the protoplasts using TRI Reagent according to the manufacturer’s protocol (Molecular Research Center,Inc). Libraries were sequenced on an Illumina NovaSeq 6000 platform using a SP 100 kit, and analyzed using the UTAP pipeline (Kohen et al. 2019) with alignment to the M82 genome (Alonge et al. 2020).
6. ATAC-seq ATAC-seq was performed on freshly isolated protoplasts. Approximately 5 mL of protoplasts at 1 × 10⁶ cells/mL were pelleted at 450 × g for 10 min with a soft start and resuspended in 5 mL ice-cold LB01 buffer(nuclei isolation protocol adopted from Doležel et al. 2007)). After 10 min incubation on ice with gentle mixing, nuclei were filtered through a 40 µm strainer, stained with DAPI (final concentration 1 µg/mL), and kept on ice in the dark until sorting. Nuclei were sorted on a BD FACSAria III equipped with a 100 µm nozzle at low speed, and ∼100,000 nuclei were collected per sample into PBS. Sorted nuclei were pelleted (500 × g, 15 min, 4 °C) and resuspended in a 50 µL tagmentation reaction containing 25 µL TD buffer, 2.5 µL TDE1 transposase, and water adjusted to volume. Tagmentation was carried out at 37 °C for 30 min. DNA was purified with the NEB Monarch PCR & DNA Cleanup Kit, eluted in 15 µL pre-warmed buffer, and quantified by Qubit. Libraries were generated using a two-step PCR amplification with KAPA HiFi polymerase and Nextera primers. The first PCR was performed for 9 cycles, followed by double-sided SPRI cleanup. A second PCR (7 cycles) was performed with the same primer pair, followed by 2× SPRI cleanup. Libraries were quantified by Qubit, assessed on a TapeStation, and sequenced on an Illumina platform using paired-end reads. Raw paired-end reads were trimmed with Trimmomatic and aligned to the *S. lycopersicum* M82 reference genome using BWA-MEM. Duplicates were removed with Picard MarkDuplicates, and reads mapping to unplaced scaffolds (chr00) were filtered out. Peaks were called with Genrich, and sgRNA-level accessibility was quantified by intersecting peak calls with guide coordinates using custom Python scripts.
7. Analysis **Custom amplicon-analysis pipeline.** Merged paired-end reads were processed with a custom Python pipeline (GitHub: https://github.com/amitcu/CRISPRIL). For each target, a reference amplicon window centered on the Cas9 cut site was extracted and indexed using 12-bp flanks. Reads containing both flanks were captured to form a fixed-length sequence window and classified as wild type or indel according to their length. This window was aligned to the amplicon reference window sequence to determine the mutation type. The pipeline outputs per-guide tables of mutation counts and unique indel events, which were used for downstream normalization and analysis.
8. **Batch effect normalization.** To control for batch effect in editing efficiency. we randomly sampled 12 guides from each experimental batch and tested them together in a single normalization batch. Editing efficiencies measured in this normalization batch were used to compute a normalization factor for each experimental batch i, using the following formula: 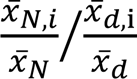 (d – original batch data, N – normalization data, i – batch). The normalized editing efficiency for each sgRNA was then obtained by multiplying its raw efficiency by the corresponding normalization factor.

## Acknowledgements

To the Israeli Ministry of Innovation, CRISPR-IL consortium to AAL for the financial support. We would like to thank Maayan Guetta, Ilan Hadad, and Ido Sela for their technical assistance, as well as all members of the Levy lab for their fruitful discussions.

## Author contributions

This work was supervised by AL. DBT, AL, AH, and AC conceptualized the study design. sgRNA selection and design, as well as calibration of the transformation protocol, were performed by AH and DBT. Protoplast isolation was done by BC, DBT, and AC. Protoplast transformation and DNA extraction were done by DBT, AH, and AC. Designing of the Illumina amplicon sequencing method, by CMB. Amplicon library preparation was done by CMB, DBT, and AC. RNA and ATAC-seq sample collection and processing were done by AC and CMB. Sequencing runs were carried out and managed by CMB. All bioinformatics and statistical analyses were carried out by AC. The manuscript was written by AC, DBT, and AL.

---

**Supplementary Figure 1.**
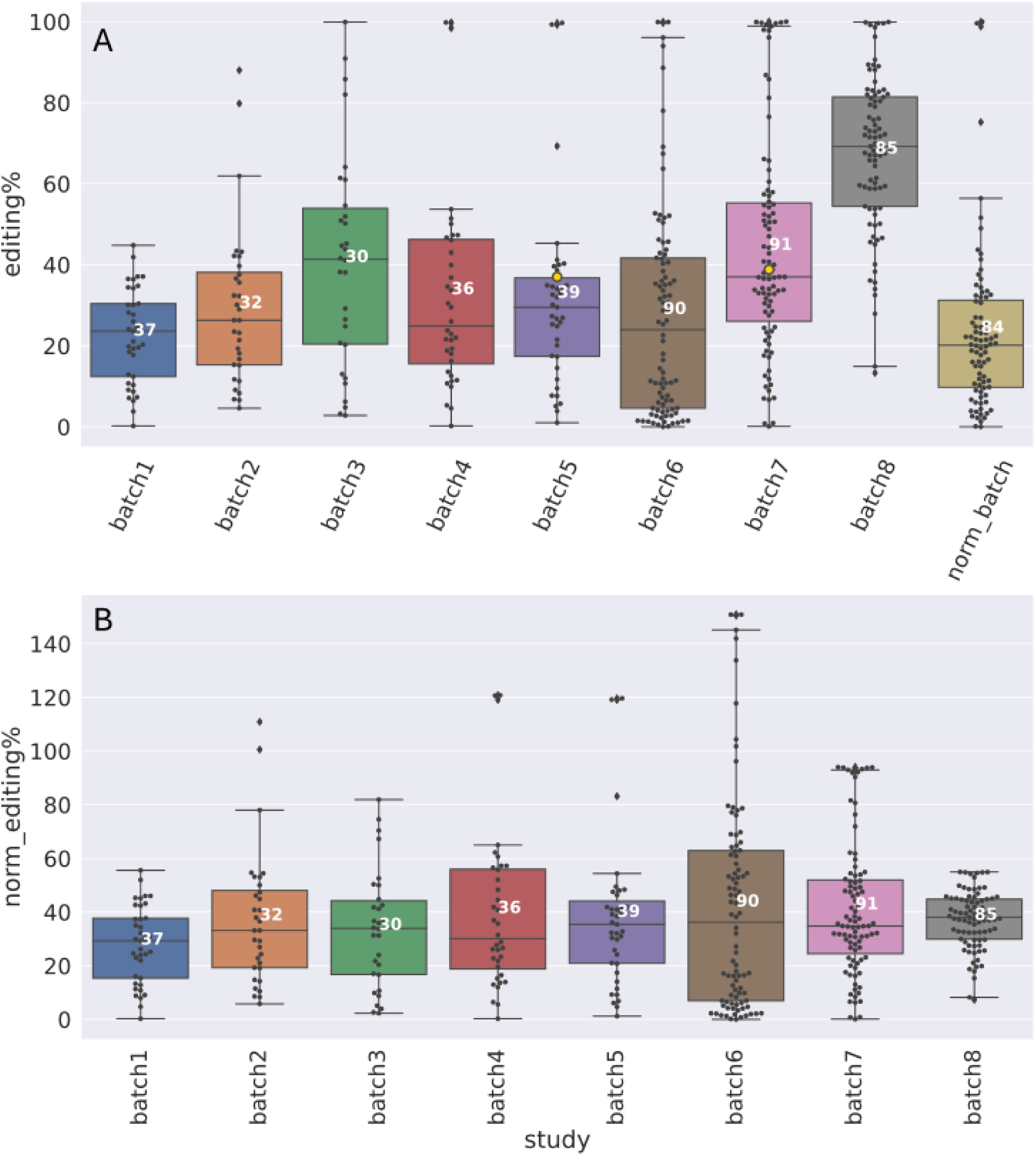
Normalization to Mitigate Batch Effects on Editing Efficiency. (A) Box plot showing the distribution of editing efficiencies across experimental batches prior to normalization. Each dot represents a single sgRNA, and white numbers indicate the number of guides tested in each batch. The "norm_batch" group contains approximately 12 sgRNAs randomly selected from each batch and processed together in a dedicated normalization run. These shared controls were used to calculate normalization factors for each batch. (B) Box plot of editing efficiencies following normalization. Editing values from each batch were adjusted using the corresponding normalization factor derived from the "norm batch" controls

**Supplementary Figure 2.**
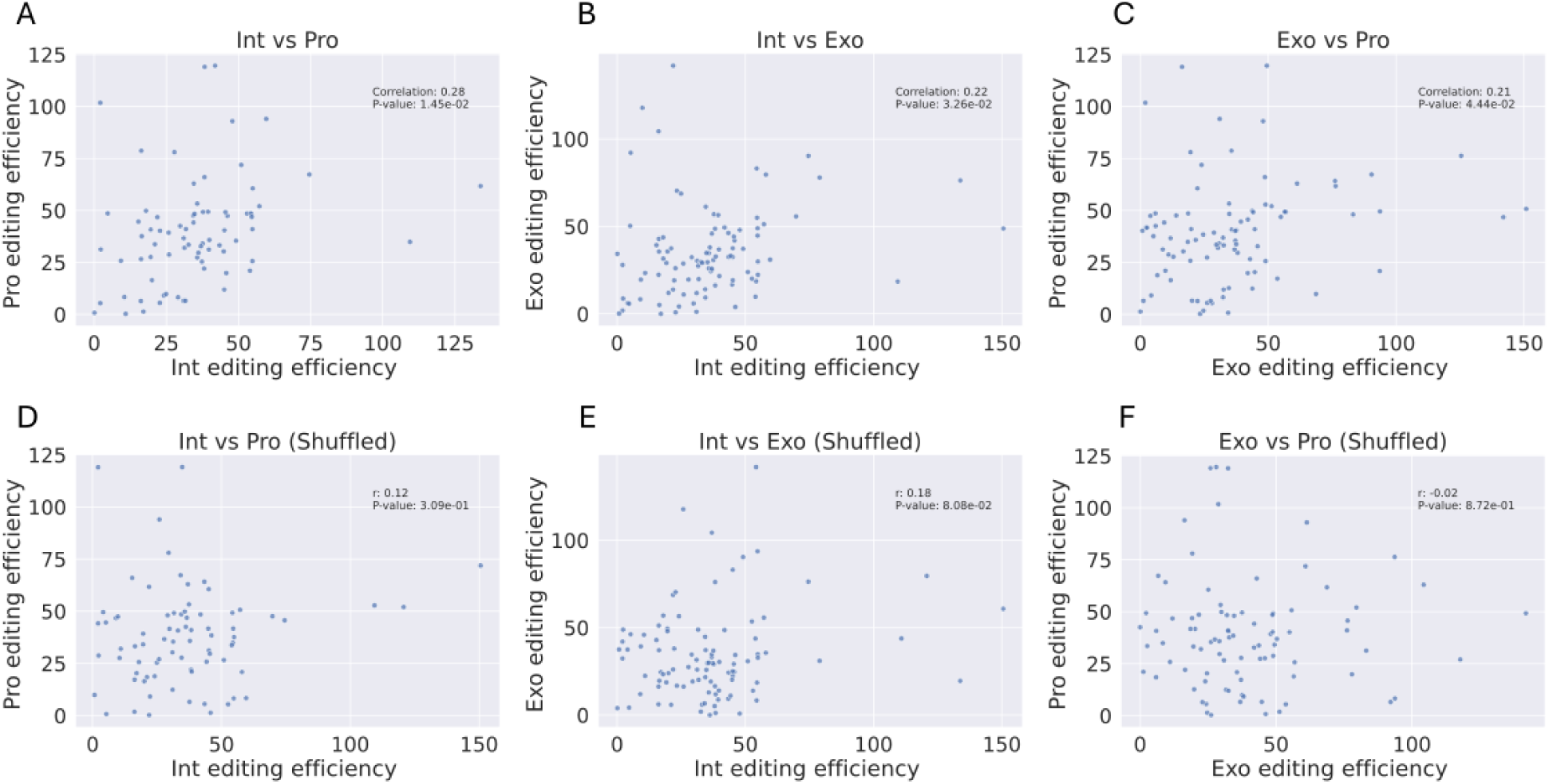
Correlation of editing efficiency between features within and between genes. Pairwise Pearson correlations of editing efficiency between sgRNAs targeting different features. **(A-C)** Correlations within the same gene: intron vs. promoter (A), intron vs. exon (B), exon vs. promoter (C). All comparisons show modest but statistically significant positive correlations (r = 0.21-0.28, p < 0.05). **(D-F)** Shuffled controls using sgRNAs targeting different genes: intron vs. promoter (D), intron vs. exon (E), exon vs. promoter (F). No significant correlations observed (r∼ 0, p > 0.5).

**Supplementary Figure 3.**
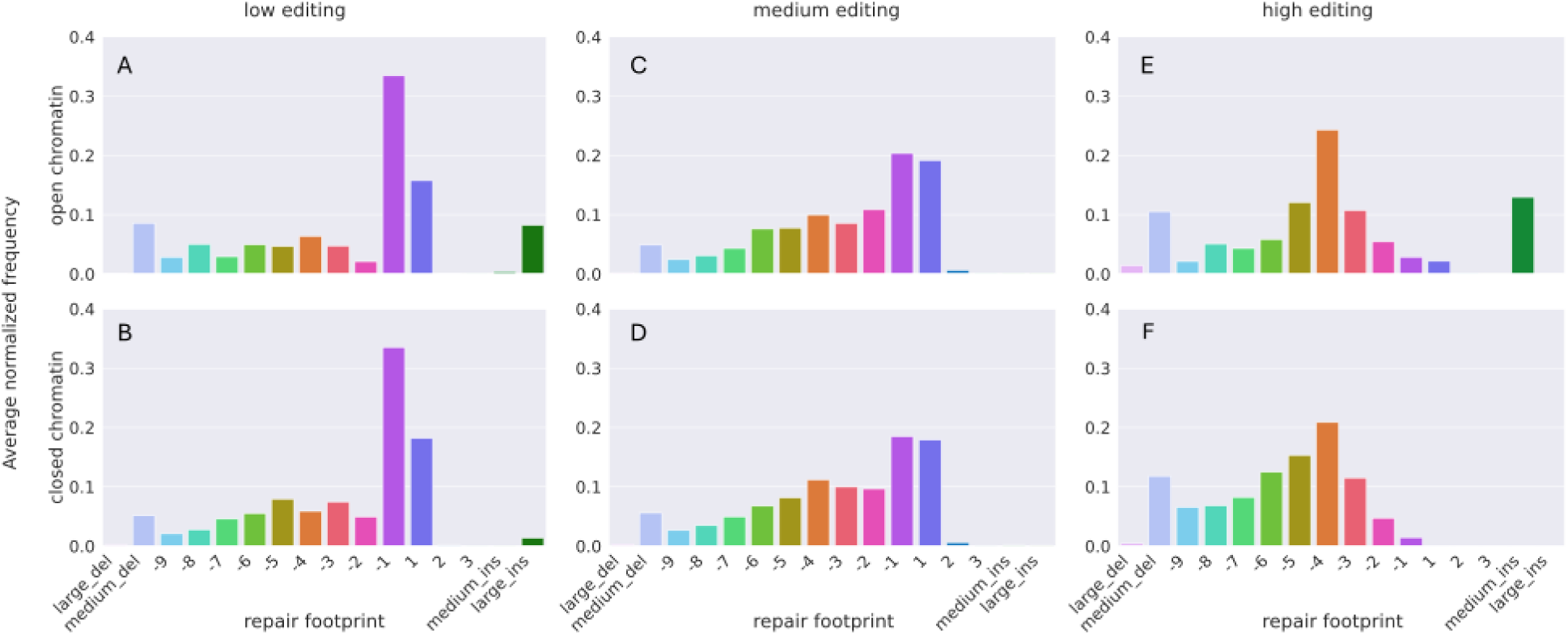
Repair footprint composition by editing efficiency and chromatin accessibility. **(A-F)** Histograms showing average repair footprint composition across editing efficiency categories, stratified by chromatin accessibility. Top panels represent open chromatin **(A, C, E);** bottom panels represent closed chromatin (**B, D, F).** Columns correspond to low **(A** (n=6), **B** (n=31)), medium **(C** (n=85), **D** (n=296)), and high **(E** (n=6), **F** (n=22)) editing efficiency groups. The x-axis indicates indel length categories, and the y-axis shows the average normalized frequency per guide.

**Supplementary Figure 4.**
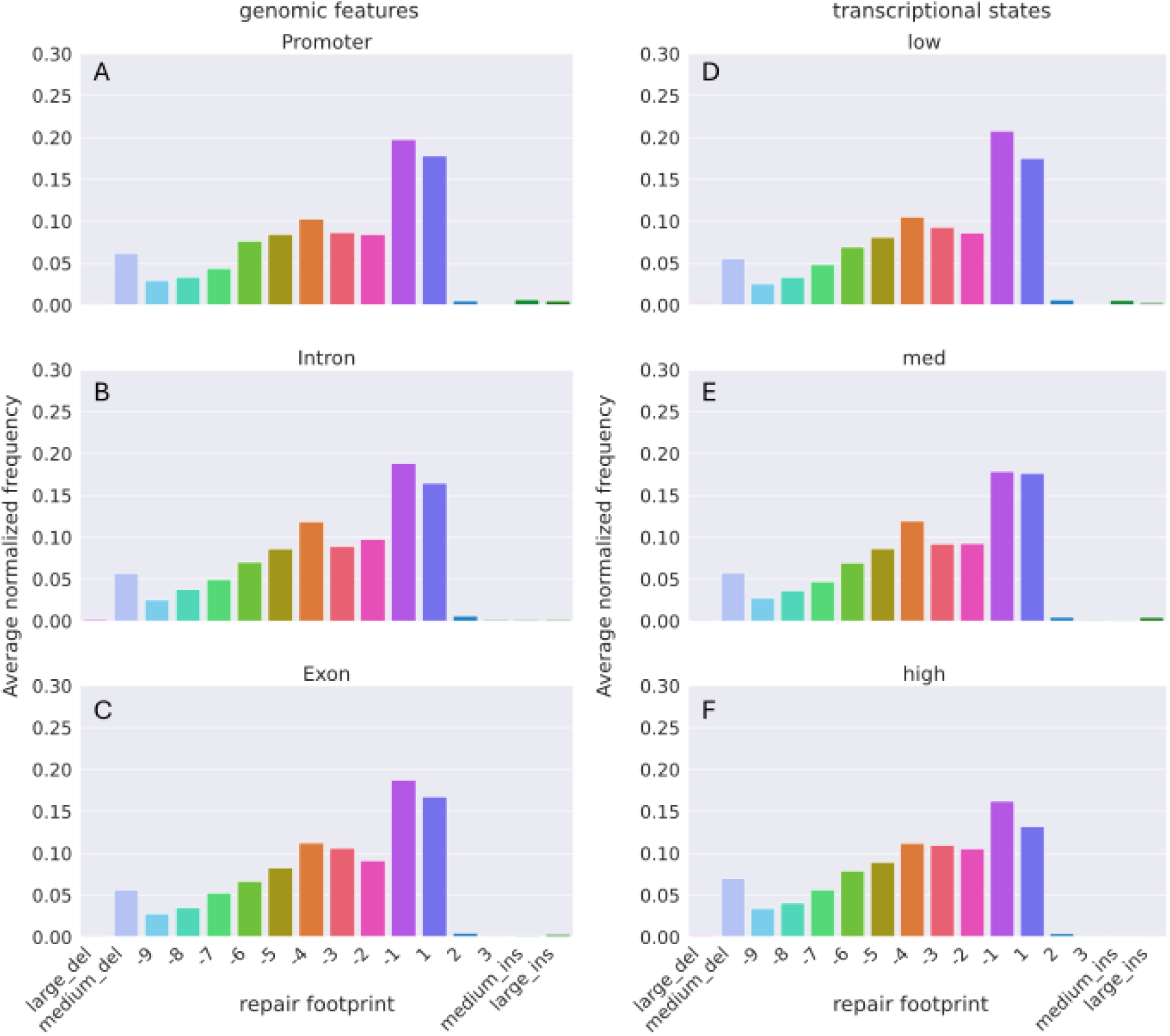
Repair footprint composition by genomic feature and transcriptional state. **(A-C)** Histograms showing average repair footprint composition for guides targeting different genomic features: exon (A), intron (B), and promoter **(C). (D-F)** Histograms showing average repair footprint composition for guides at genes with low (D), medium (E), and high (F) transcriptional states. The ×-axis indicates indel length categories, and the y-axis shows the average normalized frequency per guide. No significant differences were observed across feature or expression groups, as all distributions closely resemble the overall repair footprint profile across the dataset.

**Supplementary Figure 5.**
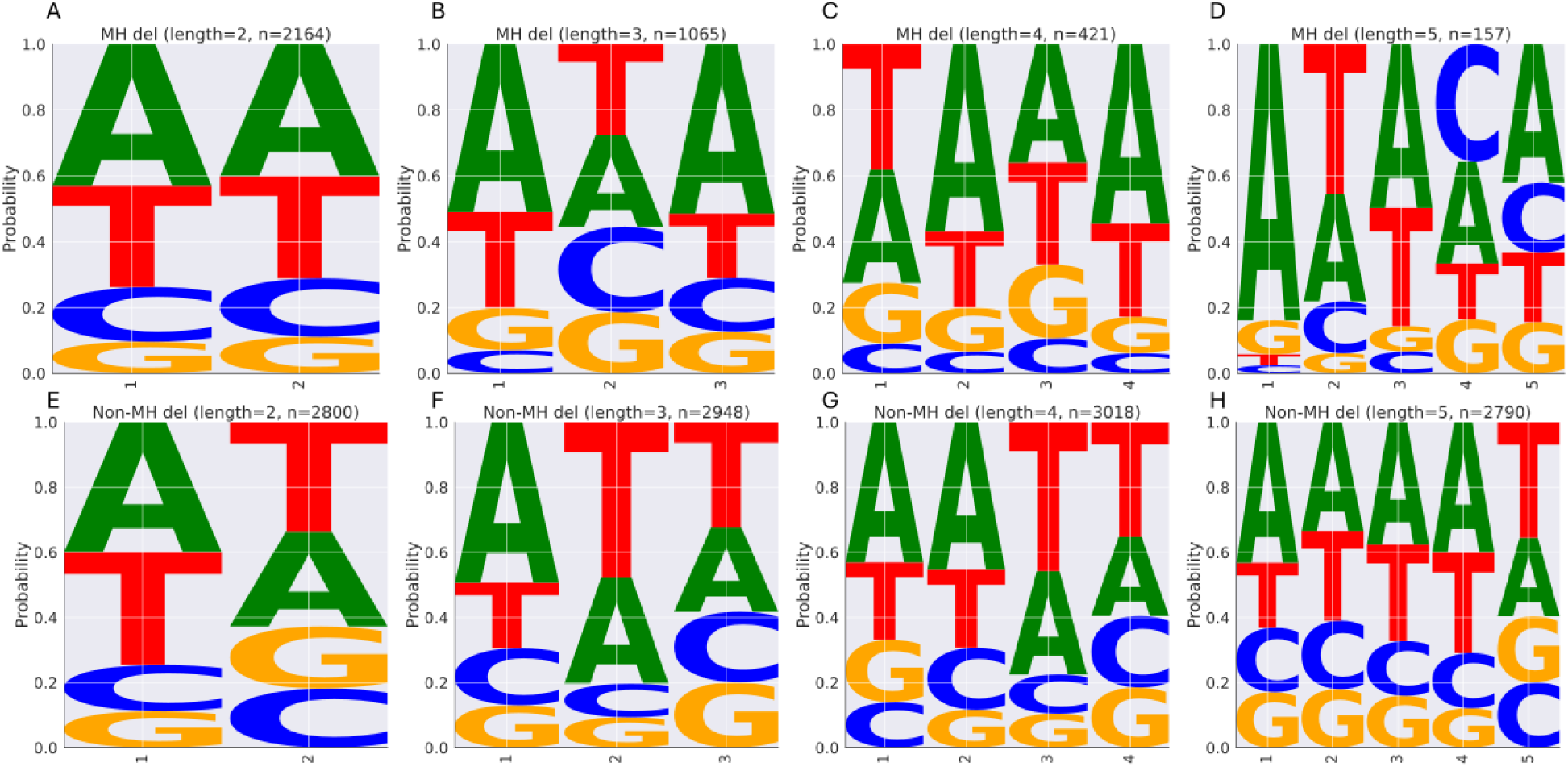
Base composition of deletion sequences stratified by deletion length and microhomology-association. Deletion base composition grouped by deletion length (2-5 bp, columns) and by association with microhomology (MH). The top row **(A-D)** shows deletions classified as MH- associated, while the bottom row **(E-H)** shows non-MH deletions. The number of deletion events contributing to each logo is indicated in the title of the panel.

